## Supplementary material for "An Open Source Unsupervised Algorithm for Identification and Fast Prediction of Behaviors": supp figs

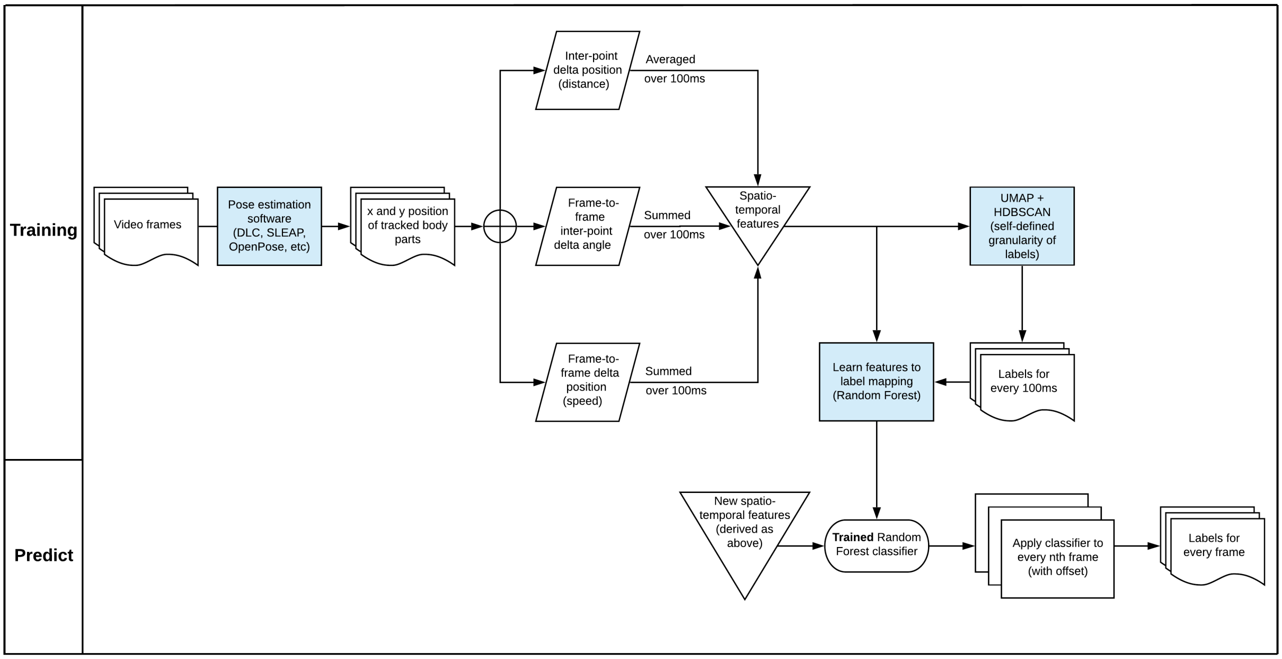


**Figure S1: B-SOiD processing diagram** A block diagram detailing the B-SOiD process flow. Video frames were filtered into pose through open source pose estimation software. Pose-relationships were then extracted from these estimates and serve as the input into unsupervised clustering (UMAP + HDB-SCAN) to assign labels. A random forest classifier is trained to learn the input pose relationships to label mapping. Once trained, the behavioral prediction based on new spatiotemporal features is much faster and can achieve higher temporal resolution of behaviors.

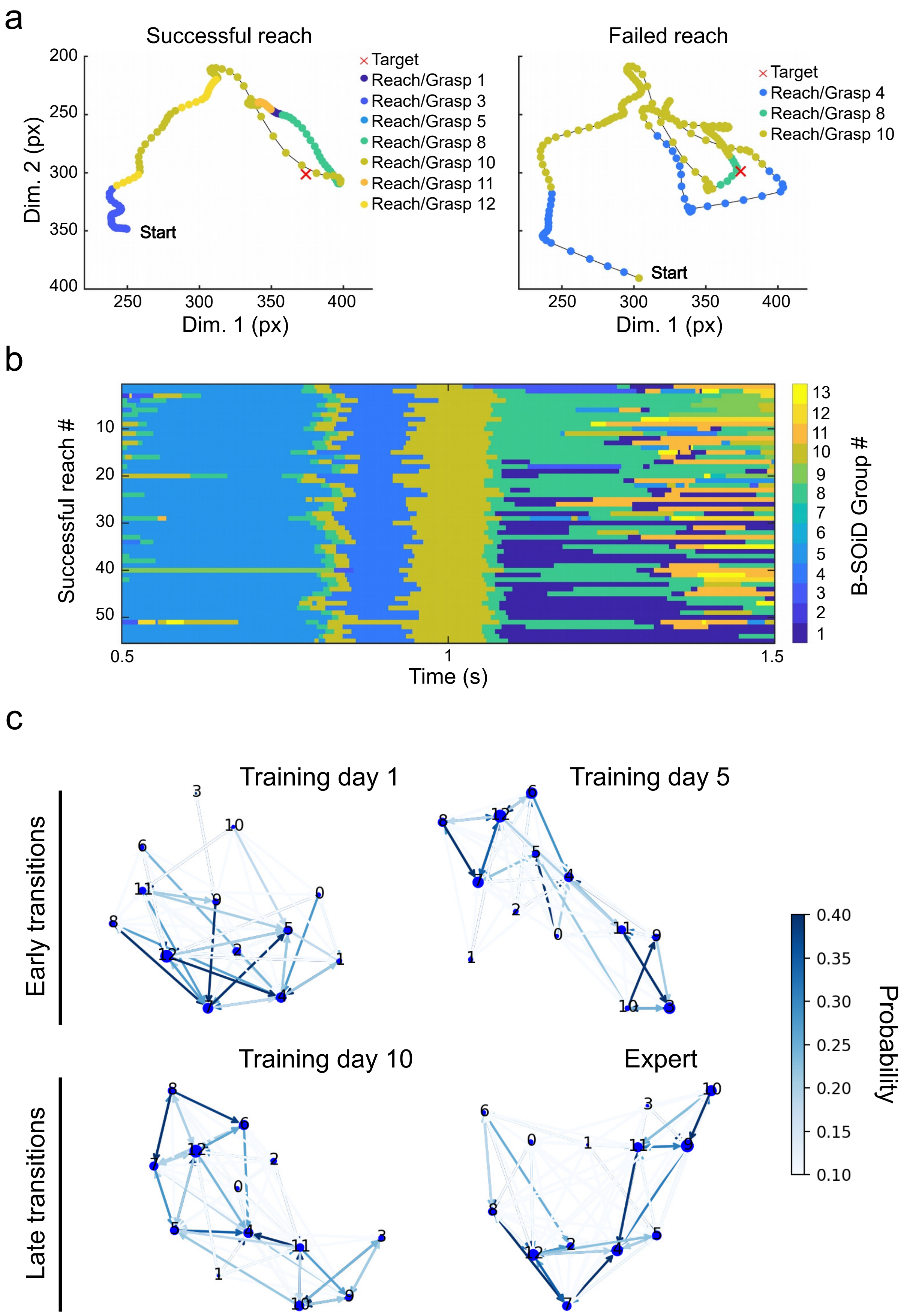


**Figure S2: Discovery and classification of sub-actions comprising rat reach and grasp task.** (a) Paw position trajectory throughout an example successful and failed attempt to reach and grasp for a pellet target. B-SOiD defined groups are mapped onto each frame of the video (300 fps taken from front, same setup as Supp Video S2 and that used in Mathis et al., 2018).

Only joint positions were used as input, and B-SOiD had no knowledge of pellet position nor whether the reach was successful or not. Note that some, but not all of the groups are shared between examples. In the fail example, the rat initiates a second swipe at the pellet, but had already knocked it off the pedestal. (b) Reach sub-action identification plotted throughout time for successful reaches across a session. Each row represents an individual reach. Time = 1s indicates detection of paw be a secondary, side view camera, which occurs just before contact with the pellet. (c) Directed graphs constructed using transition probability of these sub-actions demonstrating a more stereotypical structure in later sessions (bottom), when compared to early sessions (top). Data from the Daniel Leventhal lab.

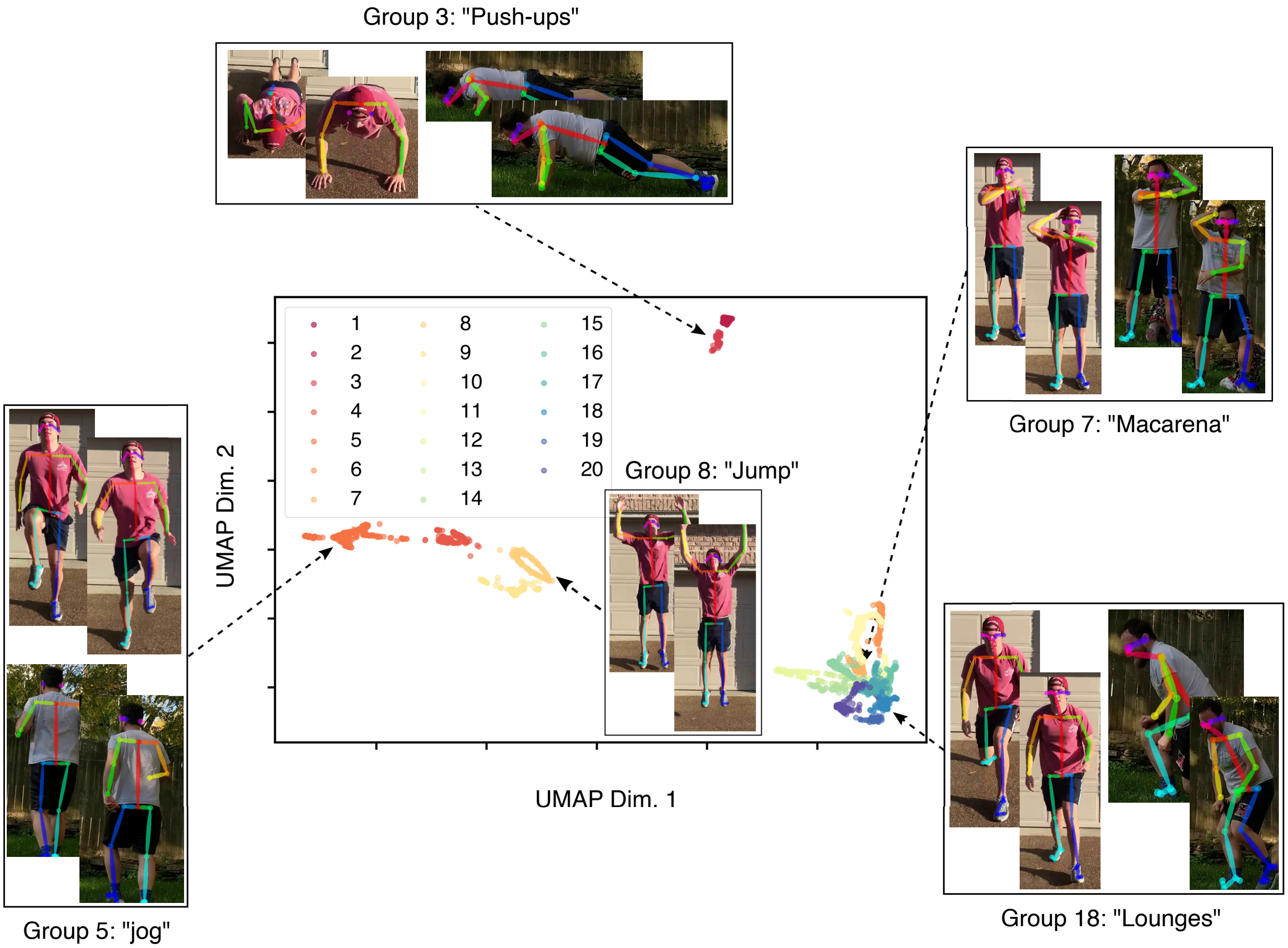


**Figure S3: B-SOiD identifies exercises and other movements from OpenPose human video data.** Video of the author working out was successfully segmented. Plot of UMAP space and representative images demonstrate that B-SOiD was able to use this human video to extract motor sub-routines. Note that regardless of individual, the same type of action occupies the same UMAP space. Rather than DeepLabCut, OpenPose software was used to provide the input pose data. Video examples, including those from a third individual, can be found in Supp Video S3.

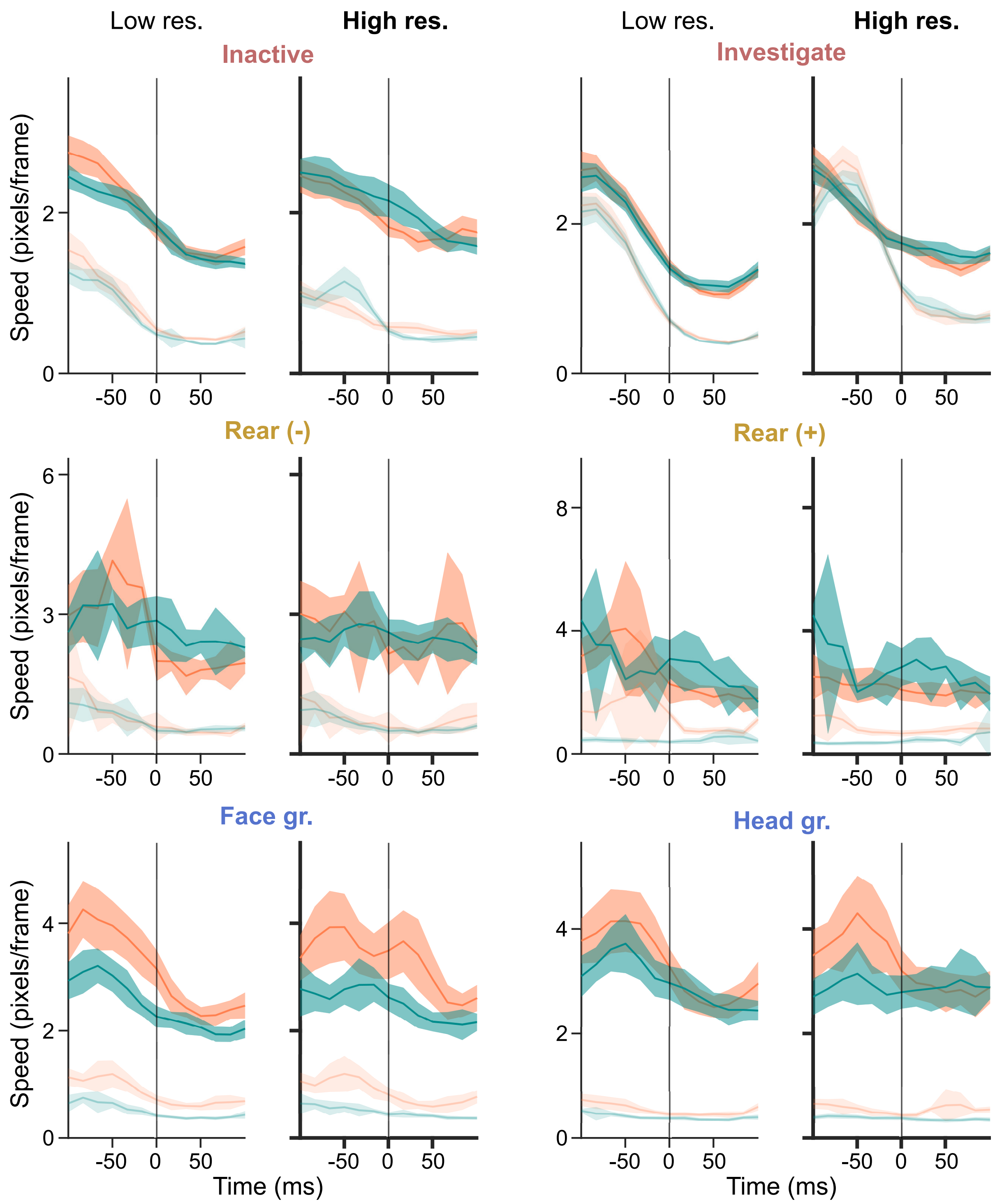

Low res.
High res.
Low res.
High res.
Inactive
Investigate
Rear (-)
Rear (+)
Face gr.
Head gr.
Speed (pixels/frame)
Time (ms)
-50
0
50

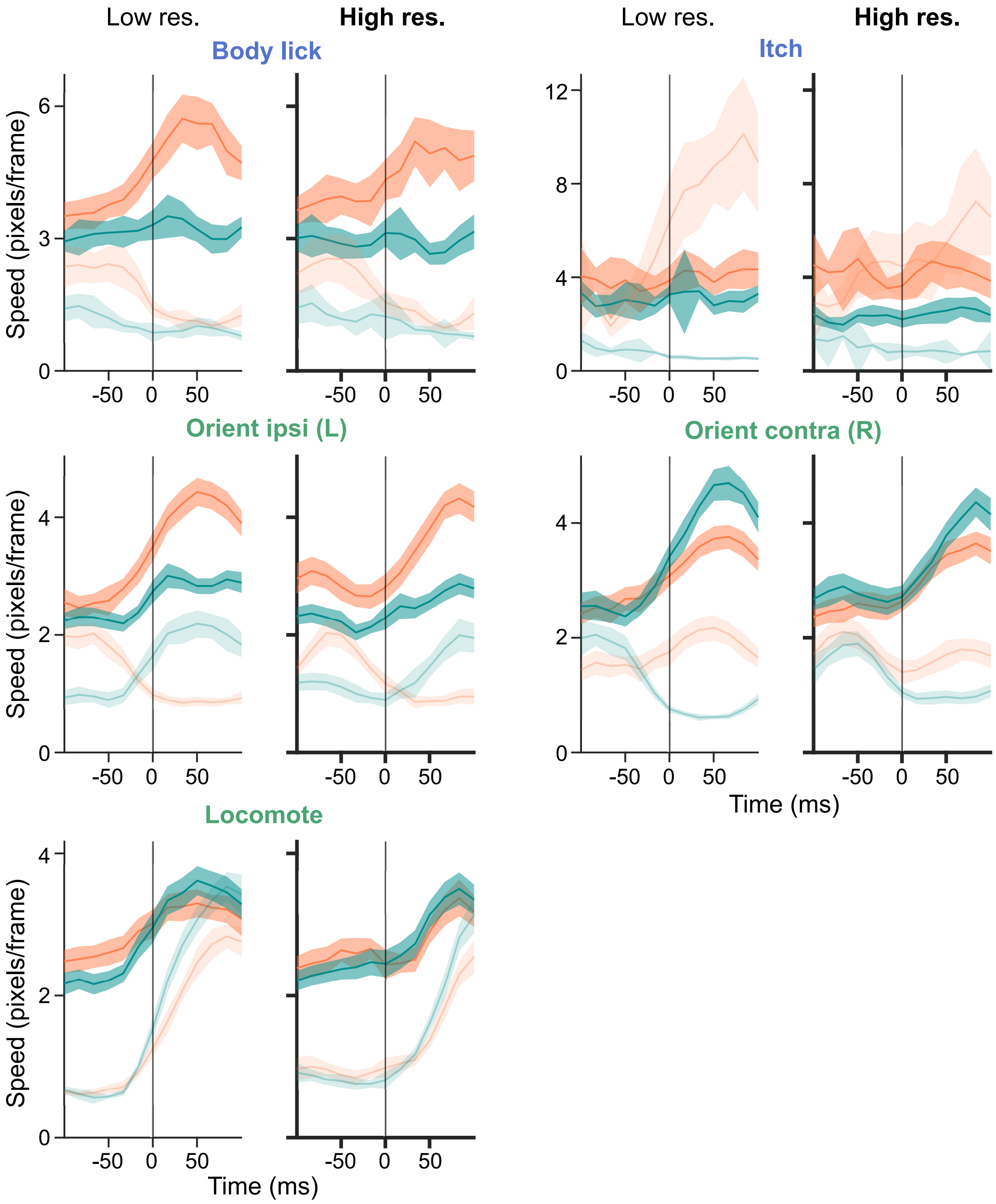


**Figure S4: Average frame-by-frame displacement aligned to movement transition time** We plotted limb speed, e.g. frame-by-frame displacement, for every transition into a given action, either for down-sampled 10fps (left, low resolution) or frameshifted 60fps (right, high resolution). Only the alignment times differ, not the frame rate frames. Colors are the same as in the individual transition plots of Fig 2d - right limbs are orange, forelimbs are darker than hindlimbs. Only +/- 100ms (one 10fps alignment) are shown in order to clarify the accuracy transition dynamics. Note the sharp inflections in average speed aligned to onset at 60fps (High res.) which is often temporally smeared at 10fps, creating an offset and smoothed transitions (Low res.).

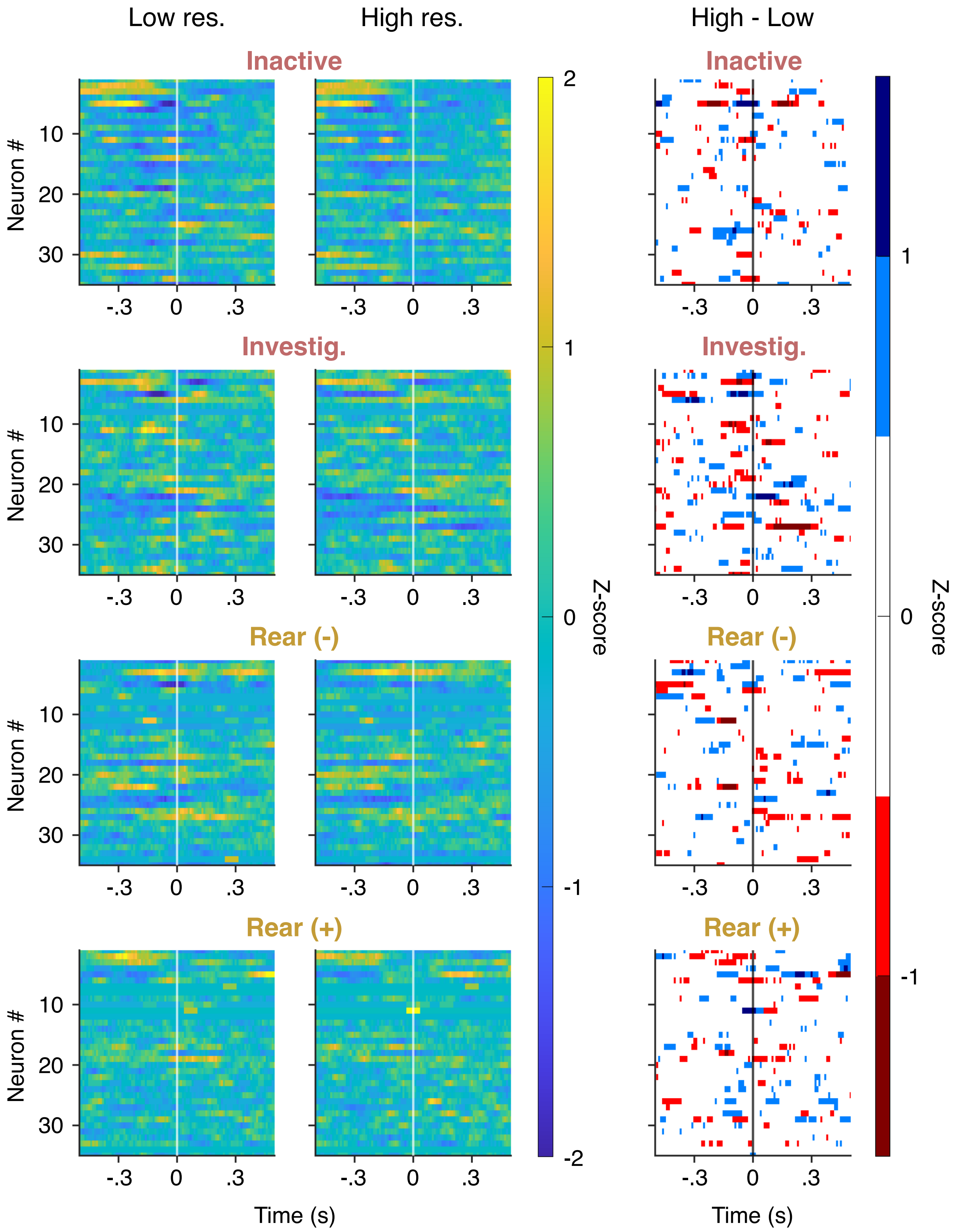

Low res.
High res.
High - Low
Inactive
Investig.
Rear (-)
Rear (+)
Neuron #
10
20
30
-.3
0
.3
Time (s)
Z-score
2
1
0
-1
-2

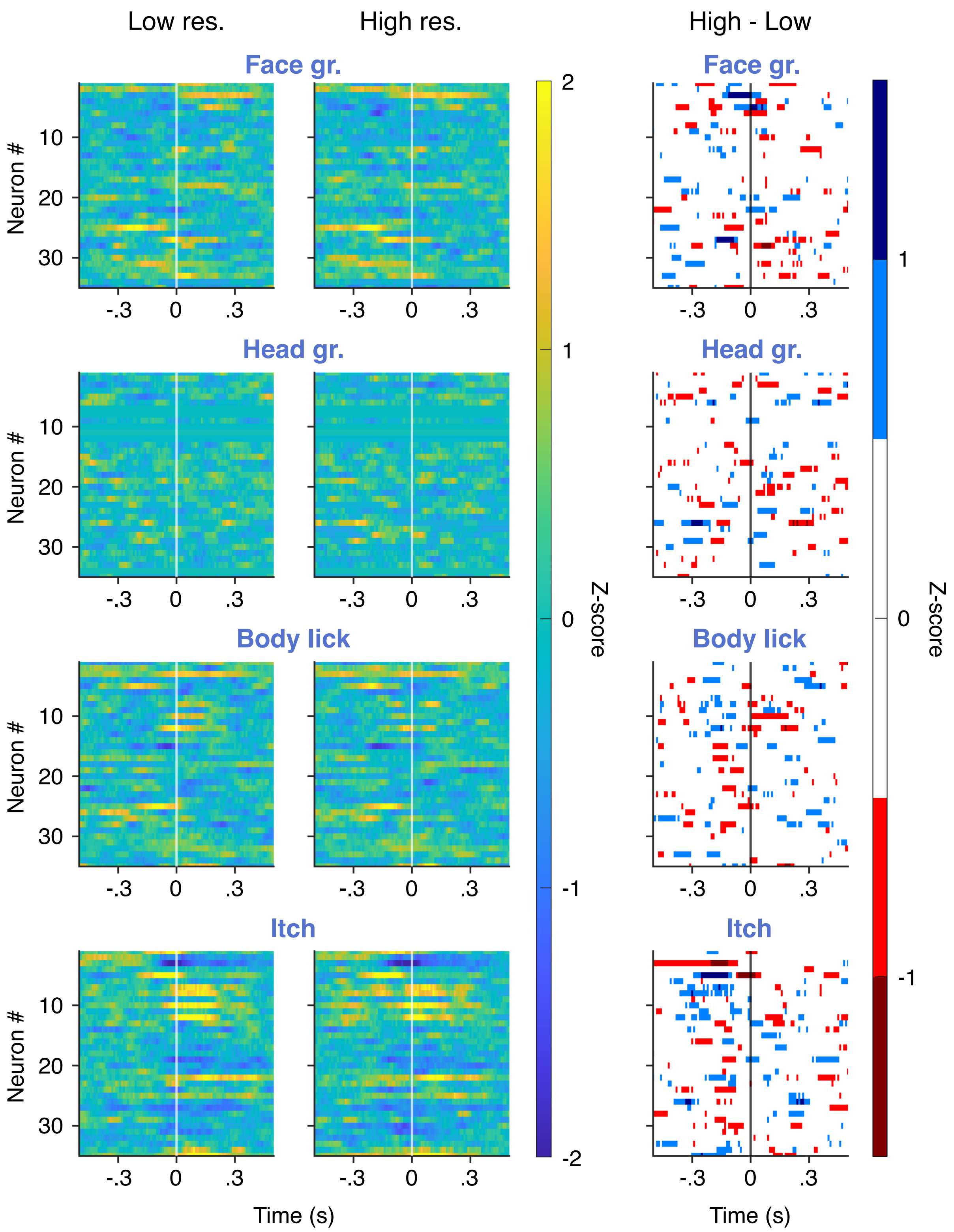

Low res.
High res.
High - Low
Face gr.
Head gr.
Body lick
Itch
Neuron #
10
20
30
-.3
0
.3
Time (s)
Z-score
2
1
0
-1
-2

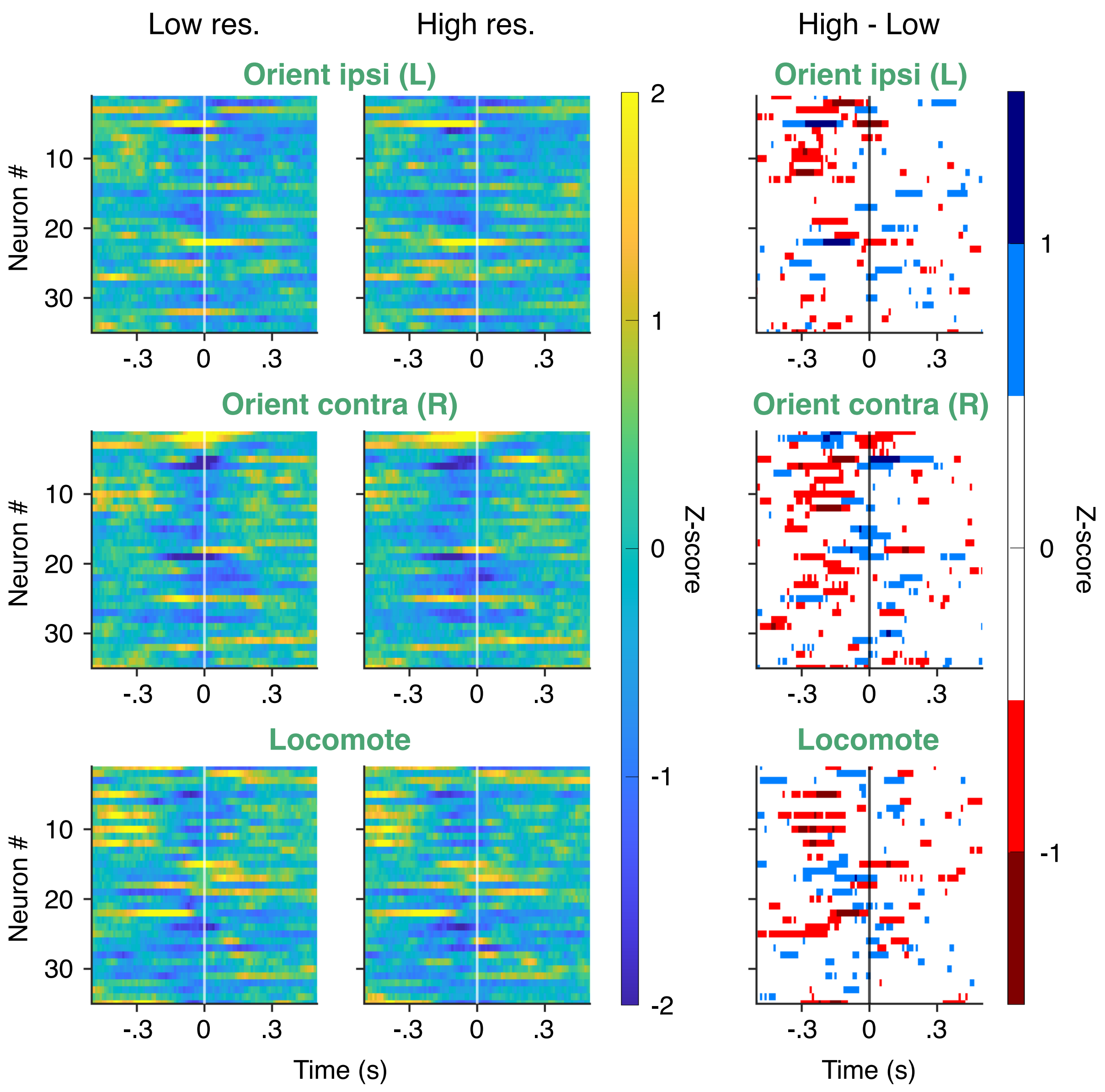


**Figure S5: Detailed neural activity for each extracted behavior.** In each set of panels, left is the observed neural response with low temporal resolution classification, middle is with high temporal resolution classification, and right is the difference between the two (high minus low resolution). Blue is indicative of a stronger signal for high resolution alignment. Neuron order is the same throughout panels, so as to allow comparison both across extraction resolutions and behaviors. Z-scores are adjusted to, and consistent for, each behavior. Behavior ID's are the same as throughout manuscript. See Supplemental Video S1 for semantic description of these behaviors.

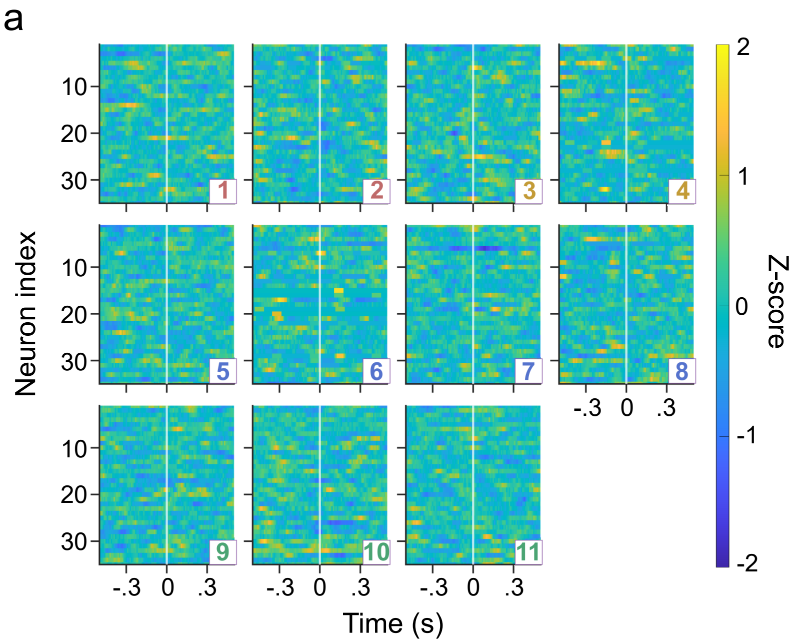


b

| Action type | Difference from scrambled data (p-val) |
|---|---|
| 1) Inactive | 3.7e-4 |
| 2) Investigate | 6.4e-7 |
| 3) Rear (-) | < 1e-7 |
| 4) Rear (+) | < 1e-7 |
| 5) Face groom | 0.026 |
| 6) Head groom | < 1e-7 |
| 7) Body lick | 0.26 |
| 8) Itch | < 1e-7 |
| 9) Orient left (ipsilateral) | < 1e-7 |
| 10) Orient right (contralateral) | < 1e-7 |
| 11) Locomote | < 1e-7 |

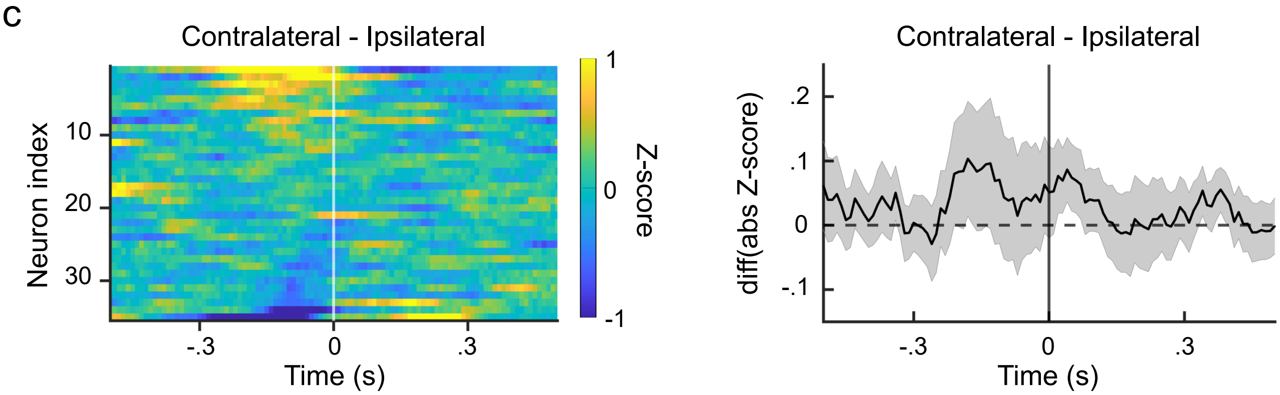


**Figure S6: Peri-event time histograms with B-SOiD compared to random alignment times.** (a) To contextualize the quality of B-SOiD-identified start times, we compared the behavior-related neural modulation relative to B-SOiD and scrambled alignment times. For each PETH, both the neuron order on the y-axis and number of alignments per behavioral group are kept the same as in Fig 3. (b) We performed Kolmogorov-Smirnov test on the magnitude z-scored activity of each neuron using frameshifted or scrambled data (period of behavior-related modulation = $\pm$ 300ms). Actions involving movements of the forelimbs produced the strongest modulation changes relative to shuffled alignments, while those actions involving less robust movements yielded smaller but still significantly larger modulations (investigate, face groom, inactive). As noted in the text, while this summary largely fits with the area recorded (caudal forelimb area of M1), it is incomplete. For instance, the only non-significant action type is ’body lick’. Forelimb movement does not play an obvious role in this behavior, but the same can be said of highly significant actions such as inactive. (c, left) Histogram of per-neuron activity for contralateral minus ipsilateral orienting behaviors. X and and Y axes the same as in Fig S3. Brighter yellow indicates greater absolute modulation for contralateral orienting. (c, right) Mean and SEM of the differences observed across the population in (left). We show these data as a glimpse of potential applications for the tool, rather than an attempt to state anything definitively about the function of motor cortex.

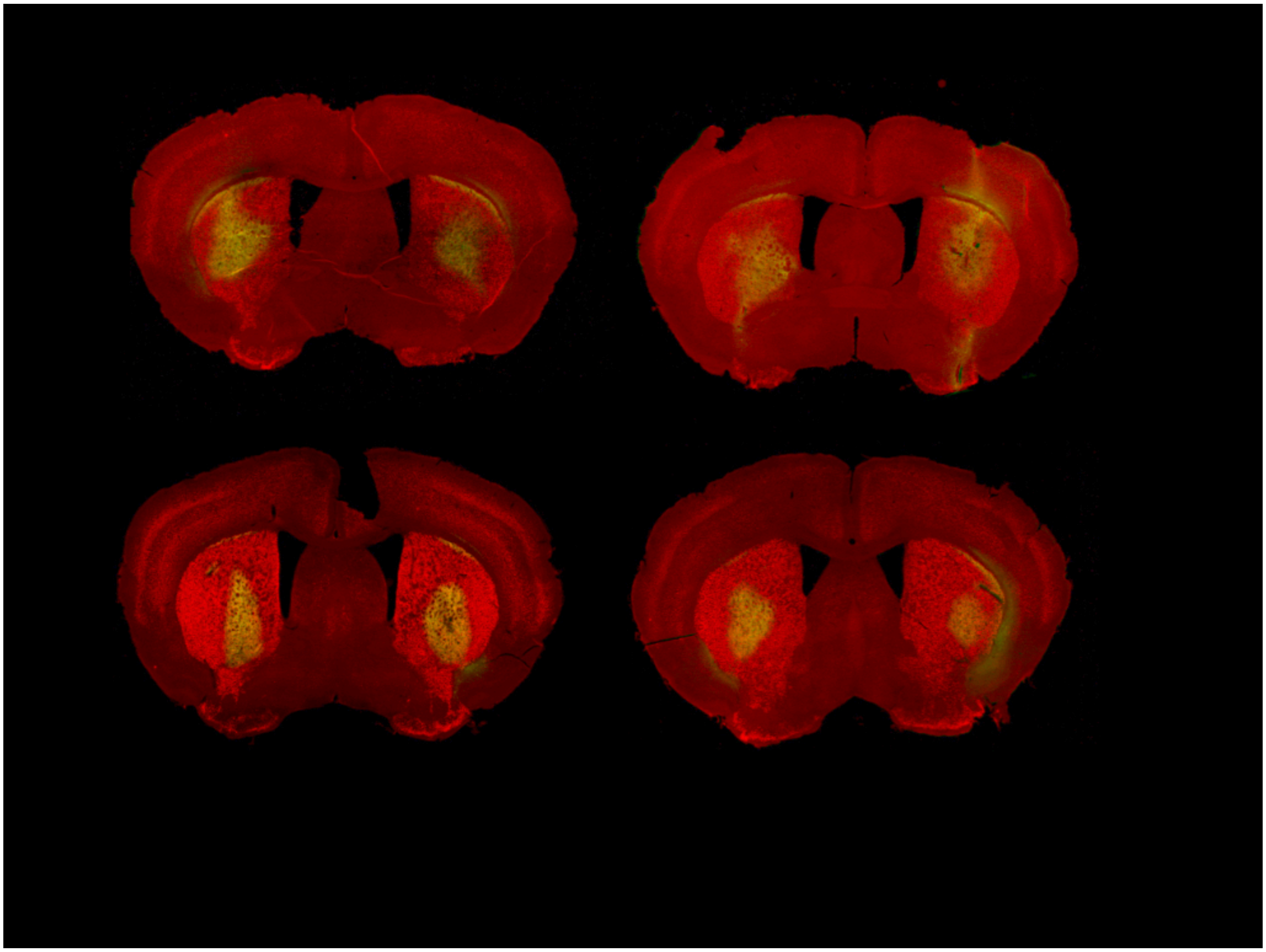

**Figure S7: Histological verification of striatal virus injection.** Coronal slices from the brains of four mice, demonstrating the spread of the GFP virus (AAV2-CAG-GFP, green) co-injected with AAV2-flex-taCasp3-TEVP. The caspase virus is designed to kill neurons in a cre-dependent manner - in this case the indirect pathway neurons in the striatum of an A2A-cre mouse. As these cells are dead (injections are made >2 weeks before experiment), the non-cre-dependent expression of GFP is used as a proxy for injection location and spread. The red background is a Foxp1 stain (overlap = yellow). Expression of GFP is restricted to the striatum. Mice provided by Mary Cundiff in the Aryn Gittis lab at Carnegie Mellon University.

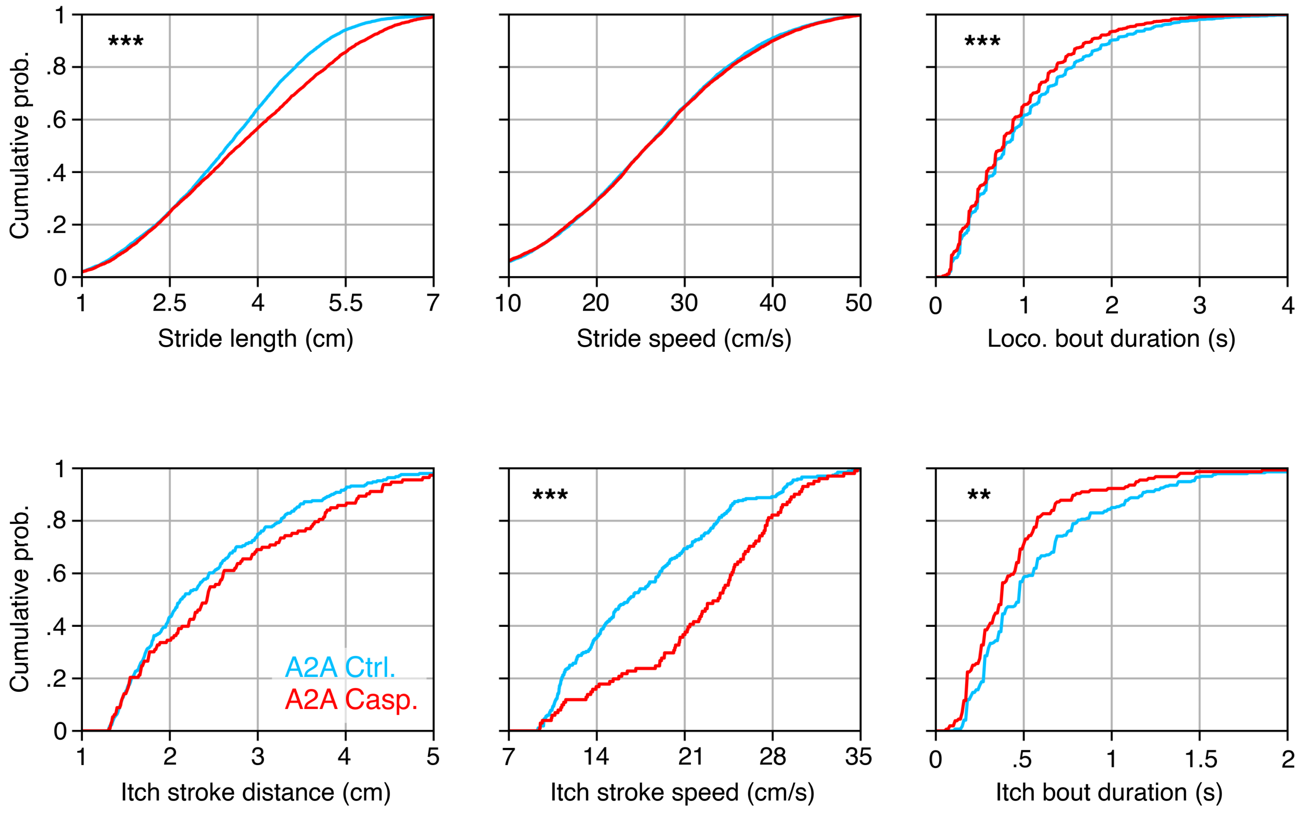


**Figure S8: Additional kinematic differences between A2A-cre control and lesioned mice.** Plots are same as Fig 6, but for locomotor (top) and itch (bottom) bouts.

Supplemental video S1: Example video demonstrating extraction of behaviors based upon natural statistical patterns of the six identified body parts. Identified frames are plotted in UMAP space as they occur. A general overview of the B-SOiD identified groups is as follows (Note that the sorted parameters were achieved without supervision by B-SOiD. No instruction as to whether a behavior should exist nor what it should look like were provided). 1) Inactive - minimal displacement speed observed at any point. 2) Investigate - minimal displacement of the hindpaws as in Inactive, but accompanied be greatly increased snout displacement and distance. 3) Rear(-) - hindpaws stationary while forepaw to hindpaw distance increased compared to inactive. Snout is partially occluded. 4) Rear(+) - hindpaws stationary while forepaw to hindpaw distance decreased compared to inactive. Snout is completely occluded. 5) Paw/Face groom - minimal distance between snout and forepaws while movements of forepaws are greater than that of snout. 6) Head groom - minimal distance between snout and forepaws while there is a smaller change in distance between forepaws and hindpaws than paw/face groom. Hindpaw to tail-base distance is reduced from paw/face groom. 7) Body lick - distance between forepaw to hindpaw is greatly reduced. Forepaws are partially occluded, whereas snout is completely occluded. 8) Itch - distance between one hindpaw and snout greatly reduced while the displacement of this same paw is greatly increased. 9) Orient left - strongly negatively skewed angle between all body points, accompanied by an increased displacement of all points. 10) Orient right - strongly positively skewed angle between all body points, accompanied by an increased displacement of all points. 11) Locomote - increased displacement of all points, and a broad distribution of distances between fore- and hind-paws on the same side (e.g. points 2 and 4, 3 and 5). No large divergence of angles observed.

Supplemental video S2: Two examples of a classified reaching sub-action in a rat. On the left, the rat commits an error in grasping the pellet due to excessive displacement between its third and fourth digits. On the right, B-SOiD identified a kinematically similar sub-action in the absence of a pellet (none was presented in this example).

Supplemental video S3: Four examples of a classified human kinesthetics across three individuals, all using the same B-SOiD model. Note the variation in performance, as well as differences in background/camera distance, cell phone camera resolution, and occasional interference. Like Supp. Video S1, the right panel of each video demonstrates the projection of that frame into UMAP space. Pose estimation achieved via OpenPose.

Supplemental video S4: The behaviors of three flies are demonstrated here. Note the consistency of action location across the UMAP parameter space. Pose estimation achieved via LEAP (Berman et al., 2014; J. R. Soc. Interface).
